# Minipoa: A minimizer-based method for fast and memory-efficient partial order alignment

**DOI:** 10.64898/2026.02.18.706716

**Authors:** Haodong Liu, Pinglu Zhang, Yanming Wei, Qinzhong Tian, Yixiao Zhai, Quan Zou, MengTing Niu

## Abstract

Partial order alignment (POA) has emerged as a fundamental component in long-read error correction, assembly and pangenomics. However, conventional POA algorithms are limited by high time and memory requirements, making them inefficient for large-scale datasets. Here, we present minipoa, a fast and memory-efficient POA tool that incorporates seed-chain-align heuristics, adaptive or static banding strategies, and single-instruction multiple-data optimizations. Minipoa achieves up to a 5-fold speedup over abPOA, reduces memory usage by up to 16-fold, and improves correction accuracy, while maintaining strong performance on both PacBio and ONT simulated datasets, and can be readily integrated into existing long-read error correction and assembly workflows. In multiple sequence alignment datasets, minipoa demonstrates superior computational efficiency and alignment accuracy compared with all other tested tools, achieving Total Column scores up to 2.5-fold higher than MAFFT in low-similarity scenarios. Moreover, minipoa enables multiple sequence alignment of megabase-long genomes and million-sequence datasets, demonstrated by 342 *Mycobacterium tuberculosis* sequences and one million SARS-CoV-2 sequences respectively. Collectively, minipoa is well positioned to become a cornerstone in the era of large-scale pangenomics.

## 1 Introduction

Partial order alignment is first introduced by Lee et al. to solve the MSA problem[1]. In POA, multiple sequence alignment (MSA) is represented as an alignment graph, specifically, a directed acyclic graph (DAG) and sequences are iteratively aligned to the alignment graph through dynamic programming (DP). The alignment graph can output not only multiple sequence alignment formats, but also a consensus sequence by applying the heaviest bundling algorithm[2]. POA plays an important role in numerous applications such as de novo genome assembly (e.g. read error correction and consensus generation)[3–6], RNA isoform inference[2], structural variant characterization[7], and variant phasing[8] or calling[9–11].

Rapid advances in sequencing technology, particularly the proliferation of long-read data and large-scale genome sequencing projects[12], have substantially increased the scale and complexity of genomic data. This rapid growth has posed significant scalability challenges for the POA method in three critical domains, including sequencing, MSA, and pangenomics[13–15].

In sequencing, the high error rates inherent to third-generation sequencing technologies necessitate highly accurate POA methods for read error correction and consensus generation[9]. Although abPOA[13] is widely used and effective, it exhibits scalability limitations in both runtime and memory usage when processing high-depth long-read datasets. In MSA, the ever-growing volume of genomic data demands tools that can perform high-quality alignment on massive datasets, a task where traditional MSA software often struggles. While POA overcomes the “once a gap, always a gap” limitation of traditional progressive methods by representing alignments as graphs[1], its scalability remains fundamentally limited by memory bottlenecks. In pangenomics, the construction of comprehensive pangenome graphs from multiple genome assemblies relies heavily on efficient graph alignment; indeed, state-of-the-art tools like Cactus[15] and PGGB[14] already depend on POA implementations as a core engine. However, aligning long stretches of homologous sequences across genomes demands substantial computing and memory resources, pushing current POA methods to their limits.

The standard POA algorithm exhibits a time and space complexity of *O*(*n* × *m*), where *n* represents the graph size and *m* denotes the sequence length. Consequently, existing POA-based methods exhibit distinct limitations when applied to large-scale data. SPOA[6] and TSTA[16] do not employ heuristic strategy to reduce DP computations, resulting in prohibitively slow performance on long sequences. Although abPOA incorporates adaptive banding techniques to accelerate alignment, it is constrained by the memory bottleneck, which prevents it from scaling to certain large datasets. POASTA[17] achieves excellent speed and memory efficiency; however, its applicability is largely restricted to datasets with extremely high sequence similarity. Recognizing graph alignment as a cornerstone for the impending era of large-scale genomics, we identify a critical need for a next-generation POA tool that offers superior performance and scalability across these diverse downstream tasks.

In this work, we present minipoa, a fast and memory-efficient POA that incorporates seed-chain-align heuristics, adaptive or static banding strategies, and single-instruction multiple-data (SIMD) optimizations. Minipoa operates in two specialized modes targeting distinct application scenarios: sequencing mode for long-read error correction and consensus generation, and MSA mode for large-scale MSA. In sequencing mode, minipoa achieves up to a fivefold speedup over abPOA while maintaining comparable correction accuracy and substantially reducing memory consumption. In MSA mode, minipoa consistently demonstrates superior computational efficiency, particularly on low-similarity datasets where traditional tools often incur prohibitive time and memory costs. Notably, it is capable of completing certain alignment tasks that other tools fail to handle, such as aligning 342 *Mycobacterium tuberculosis* sequences (each of megabase length) or one million SARS-CoV-2 sequences. Additionally, minipoa also supports Graphical Fragment Assembly (GFA) output, facilitating seamless integration with downstream graph genomics tools and applications.

## 2 Result

Minipoa constructs a graph by incrementally integrating input sequences in FASTA format, with the option to initialize the graph from an existing GFA file. From the final graph, minipoa can generate consensus sequences, multiple sequence alignments, or export the graph in GFA format (Figure 1A). Efficient alignment is achieved through a consensus-guided strategy (Figure 1B). Minipoa is implemented in C++. It supports two distinct operational modes: sequencing and MSA, each controlled by a specific set of predefined parameters. Minipoa is designed to be user-friendly, requiring only basic parameters, with well-defined default values for more complex parameters (see Supplemental Method). All experiments were conducted on a Linux server running Ubuntu 22.04.5 LTS with AMD EPYC 7763 CPUs (256 cores, 2.45 GHz), 1 TB RAM.

**Figure 1.**
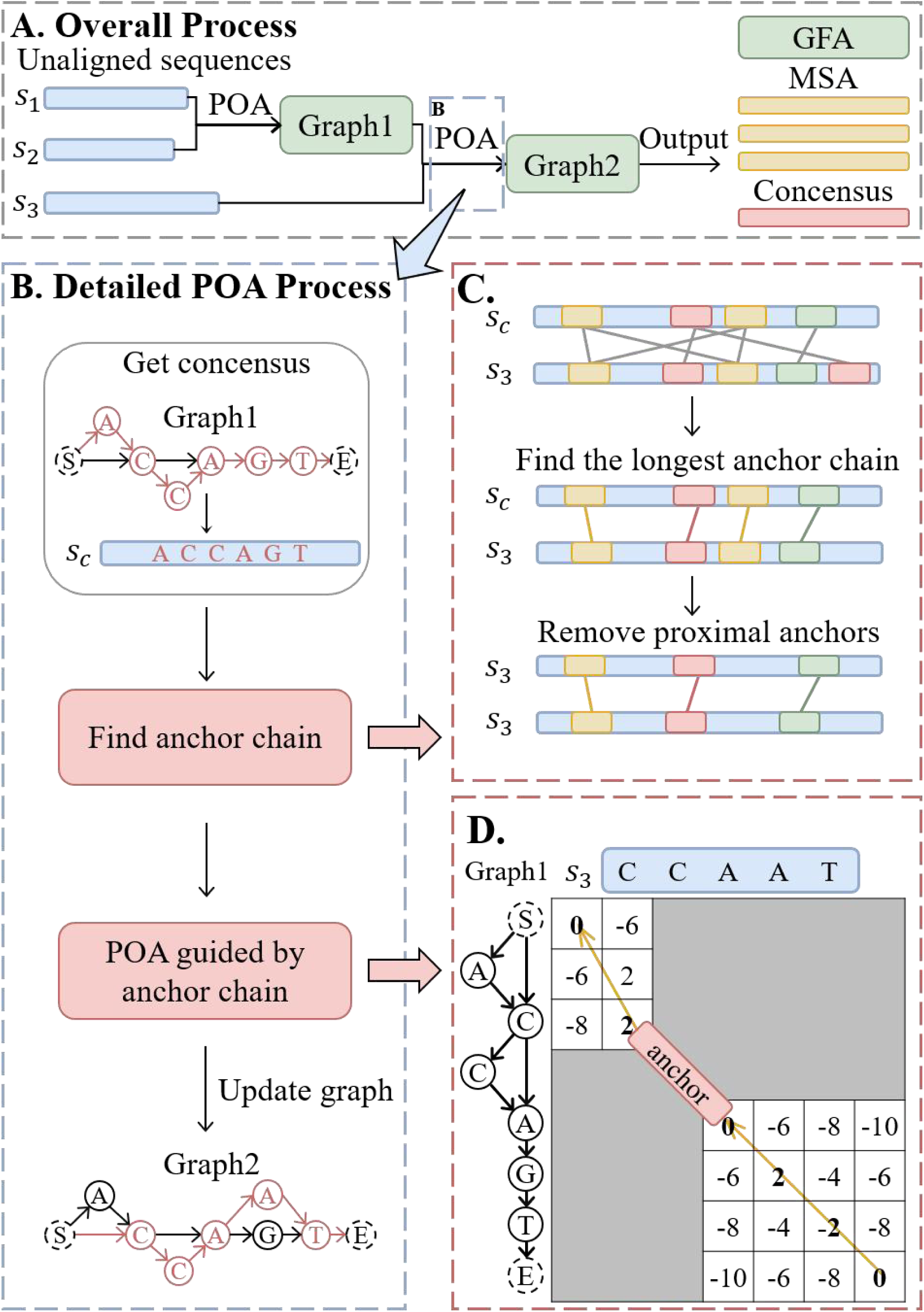
The flowchart of minipoa. (A) Overall workflow of minipoa, where unaligned sequences are iteratively aligned to update the graph, ultimately producing outputs in three formats: a consensus sequence, a multiple sequence alignment, or a GFA graph. (B) Detailed POA workflow, showing how a consensus sequence is generated from the current graph and used to guide the alignment of a query sequence. (C) The procedure for finding an anchor chain, including anchor collection, longest-chain selection, and removal of proximal anchors. (D) Schematic illustration of the core POA procedure in minipoa, where anchor chain guide the sequence-to-graph alignment.

To comprehensively evaluate the performance of minipoa, we conducted a rigorous series of experiments spanning both sequencing and MSA modes. For the sequencing mode, we assessed read error correction capabilities using simulated datasets with varying sequence lengths and sequencing depths (using single thread), and further validated practical utility by integrating minipoa into the Racon pipeline for error correction on five real-world datasets (using 16 threads). For the MSA mode, we benchmarked alignment accuracy and computational efficiency against state-of-the-art tools using simulated sequences with varying similarities (using single thread) as well as five diverse real genomic datasets (using 8 threads). Finally, to demonstrate scalability and robustness on large-scale datasets, we performed alignments on 342 Mycobacterium tuberculosis sequences or one million SARS-CoV-2 sequences (using 16 threads).

### 2.1 Evaluation metrics

For the sequencing experiments, we evaluated the quality by calculating the error rate of the generated consensus sequence. Each consensus sequence or corrected reads was aligned to its corresponding original reference sequence using Minimap2[18]. The error rate was defined following the formulation in [13], as the total number of mismatches, insertions, and deletions in the alignment divided by the length of the consensus sequence or the corrected reads.

To assess the quality of MSA, we employed several established benchmark metrics: the Sum-of-Pairs (SP) score[19], Q score, and Total Column (TC) score[20]. Adopting the scoring scheme from FMAlign[21], the SP score is computed by summing the scores of all pairwise comparisons within the alignment, where a match is assigned a score of 1, a mismatch -1, and a gap -2; thus, a higher SP score indicates better alignment accuracy. To enable fair comparison across datasets of varying lengths, we used the Scaled-SP score, which normalizes the raw SP score by the sequence length. The Q score, ranging from 0 to 1, quantifies the similarity between a test alignment and a reference alignment, with a value of 1 denoting complete agreement. The TC score is a metric that evaluates the proportion of correctly aligned columns relative to a reference alignment.

### 2.2 Experimental Results under Sequencing Mode

#### 2.2.1 Read Error Correction on Simulated Data

To simulate long-read datasets with varying read lengths and sequencing depths, we first randomly extracted a single 2 million bp continuous reference sequence from the GRCh38 human reference genome[22]. From this sequence, five subsequences with lengths of 500 bp, 5,000 bp, 20,000 bp, 50,000 bp, and 100,000 bp were further extracted and used as reference sequences for simulation. For each reference length, three sequencing depths (10×, 30×, and 50×) were considered, resulting in a total of 15 (5×3) simulation settings. For each setting, 100 independent clusters of sequences were generated from the same reference using PBSIM2[23], each representing an independent random simulation, with error profiles corresponding to Oxford Nanopore Technologies (ONT) or Pacific Biosciences (PacBio) sequencing. We evaluated minipoa on these simulated datasets and compared its performance with abPOA and TSTA (see Supplemental Method). All three tools exhibited comparable performance on both ONT and PacBio simulations (Figure 2 and Figure 3); therefore, we focus the subsequent analysis on PacBio data for clarity and consistency.

**Figure 2.**
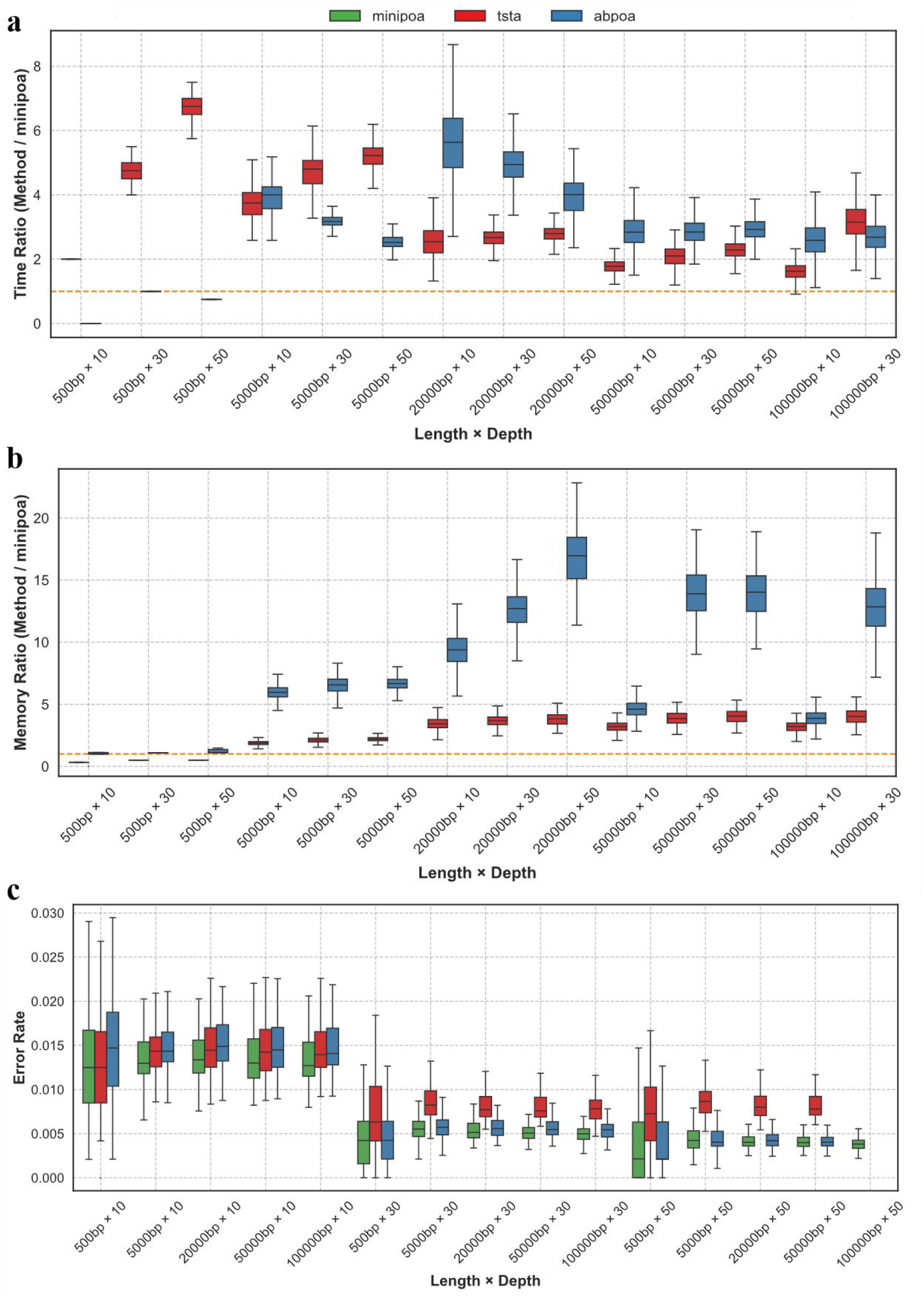
Performance comparison of minipoa, TSTA, and abPOA across different sequence lengths and sequencing depths on simulated PacBio datasets. (a) Runtime. (b) Memory usage. (c) Consensus error rate.

**Figure 3.**
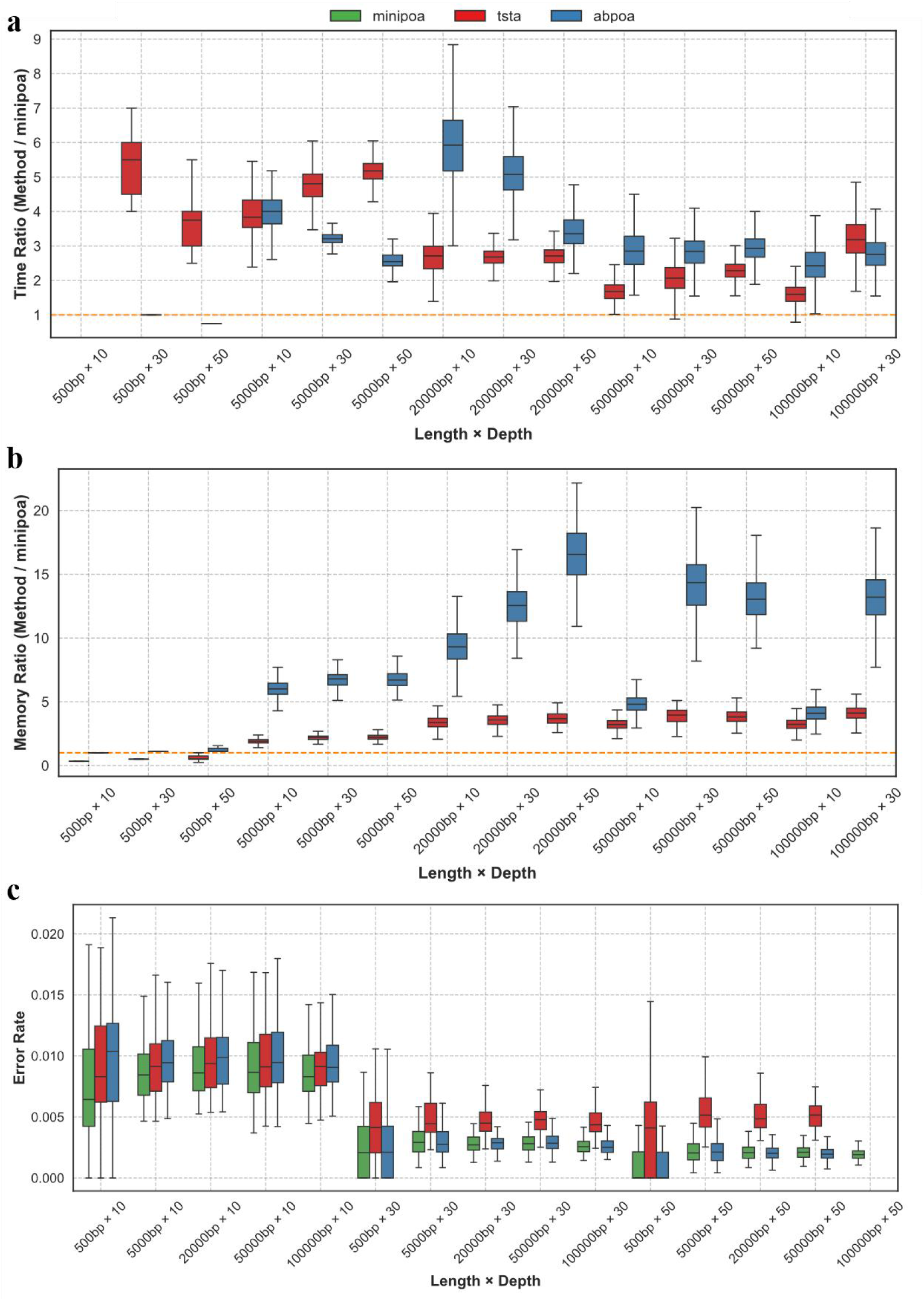
Performance comparison of minipoa, TSTA, and abPOA across different sequence lengths and sequencing depths on simulated ONT datasets. (a) Runtime. (b) Memory usage. (c) Consensus error rate.

As shown in Figure 2, minipoa demonstrates clear advantages in computational efficiency and memory usage across varying sequence lengths and sequencing depths. Specifically, minipoa is 2.5∼5.5 times faster than abPOA, excluding the small dataset with read length of 500 bp (Figure 2a). Regarding resource consumption, minipoa reduces peak memory usage by 3.8∼16.5 times relative to abPOA, again excluding the small 500 bp dataset (Figure 2b). Moreover, both abPOA and TSTA failed to complete due to memory exhaustion when processing sequences of length 100,000 at 50 × depth, whereas minipoa maintained high correction accuracy. These results highlight that, even under challenging conditions with long sequences and low coverage, minipoa demonstrates robust alignment and consensus generation, ensuring reliable performance across diverse scenarios.

In terms of correction accuracy, minipoa consistently achieved low error rates across varying sequence lengths and sequencing depths, performing comparably to or surpassing the current state-of-the-art method, abPOA. Notably, as sequencing depth increased, both minipoa and abPOA exhibited a general decrease in error rates. However, TSTA showed a rebound in error rate at 50× coverage (Figure 2c). Moreover, compared to TSTA without adaptive banding, minipoa with adaptive banding significantly reduces the run time in all simulation settings without sacrificing the alignment accuracy. In summary, these results demonstrate that minipoa not only generates high-quality consensus sequences from error-prone long reads but also provides significant improvements in speed and memory efficiency over existing state-of-the-art tools.

#### 2.2.2 Read Error Correction on Real Data

We further evaluated the use of minipoa within a popular longread error correction pipeline Racon[6] on five real datasets (see Supplemental Table S1) from different genomes, including *Lambda phage*, *E. coli K-12*, *S. cerevisiae S288c*. We integrated minipoa into Racon (Racon-minipoa) and ran it in the error correction mode on all datasets. For comparison, we also tested Racon-SPOA, which uses SPOA for consensus generation, and Racon-abPOA, which employs abPOA (see Supplemental Method).

The computational efficiency of different consensus backends integrated into Racon was first compared. As shown in Table 1, Racon-minipoa consistently required less total runtime and consensus calling time than both Racon-SPOA and Racon-abPOA across all real datasets. The runtime advantage was particularly evident on large eukaryotic datasets. For example, on the *S. cerevisiae S288c* ONT dataset, Racon-minipoa completed consensus generation in 44.66 minutes, achieving a 1.74×speedup over Racon-SPOA, while Racon-abPOA was substantially slower. These results demonstrate that minipoa can significantly improve the efficiency of long-read error correction when used as the consensus module in Racon.

**Table 1.**
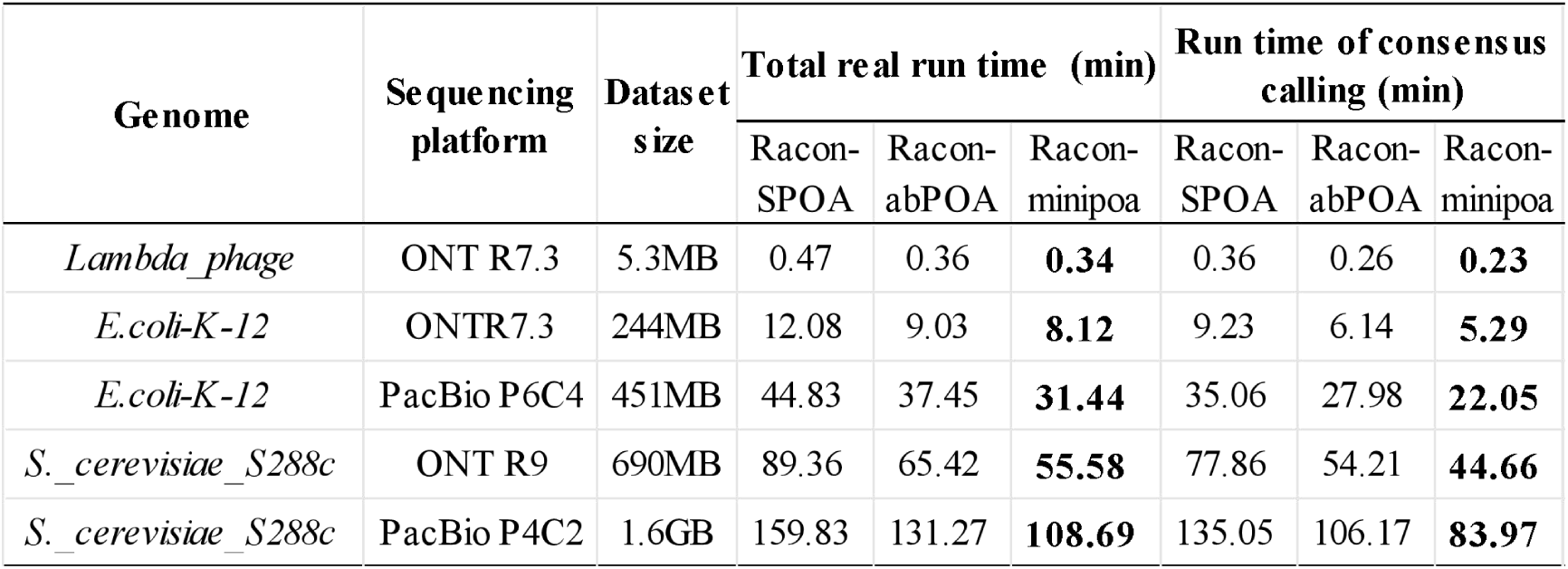
Real runtime and consensus calling time of Racon-SPOA, Racon-abPOA and Racon-minipoa on five real datasets from different genomes.

Regarding correction accuracy, Table 2 presented the error rates of mappable corrected reads. Racon-minipoa achieved the lowest error rates in most datasets, particularly demonstrating superior performance on the ONT sequencing platforms (*Lambda phage*, *E. coli K-12*, and *S. cerevisiae S288c*). Although the error rates of Racon-minipoa on the PacBio datasets (*E. coli K-12* and *S. cerevisiae S288c*) were slightly higher than those of Racon-SPOA, it remained competitive. Notably, Racon-minipoa consistently outperformed Racon-abPOA across all datasets, exhibiting significantly lower error rates in both ONT and PacBio sequencing platforms. These results demonstrate that minipoa not only accelerates consensus generation but also preserves correction accuracy.

**Table 2.** Number of mappable corrected reads, and corrected reads’ error rate of Racon-SPOA, Racon-abPOA and Racon-minipoa on five real datasets from different genomes.

| Genome | Sequencing platform | total reads | Raw error rate (%) | mappable corrected reads |  |  | Error rate of mappable corrected reads (%) |  |  |
| --- | --- | --- | --- | --- | --- | --- | --- | --- | --- |
|  |  |  |  | Racon-SPOA | Racon-abPOA | Racon-minipoa | Racon-SPOA | Racon-abPOA | Racon-minipoa |
| *Lambda phage* | ONT R7.3 | 646 | 18.20 | **515** | **515** | **515** | 3.34 | 5.86 | **3.21** |
| *E.coli-K-12* | ONTR7.3 | 25483 | 13.46 | **24164** | **24164** | **24164** | 2.16 | 4.28 | **1.81** |
| *E.coli-K-12* | PacBio P6C4 | 53332 | 12.42 | 51283 | **51286** | 51284 | **1.35** | 6.10 | 1.63 |
| *S. cerevisiae_S288c* | ONT R9 | 119955 | 13.21 | **103921** | **103921** | **103921** | 4.81 | 4.70 | **4.19** |
| *S. cerevisiae_S288c* | PacBio P4C2 | 269145 | 13.57 | 237968 | 237985 | **237990** | **3.06** | 7.91 | 3.50 |

### 2.3 Experimental Results under MSA Mode

#### 2.3.1 Multiple Sequence Alignment on Simulated Data

For the comparative evaluation of MSA quality, we simulated mitochondrial-like sequences using INDELible[24] with sequence similarities ranging from 70% to 99%, adopting the parameter settings described in FMAlign2[25]. Figure 4 presents a detailed comparison of minipoa, abPOA [13], MAFFT[26] (Auto mode), MUSCLE[20], and ClustalΩ[27] on simulated datasets, evaluating performance and accuracy in terms of scaled SP score, Q score, TC score, as well as computational time and memory consumption across different sequence similarity levels.

**Figure 4.**
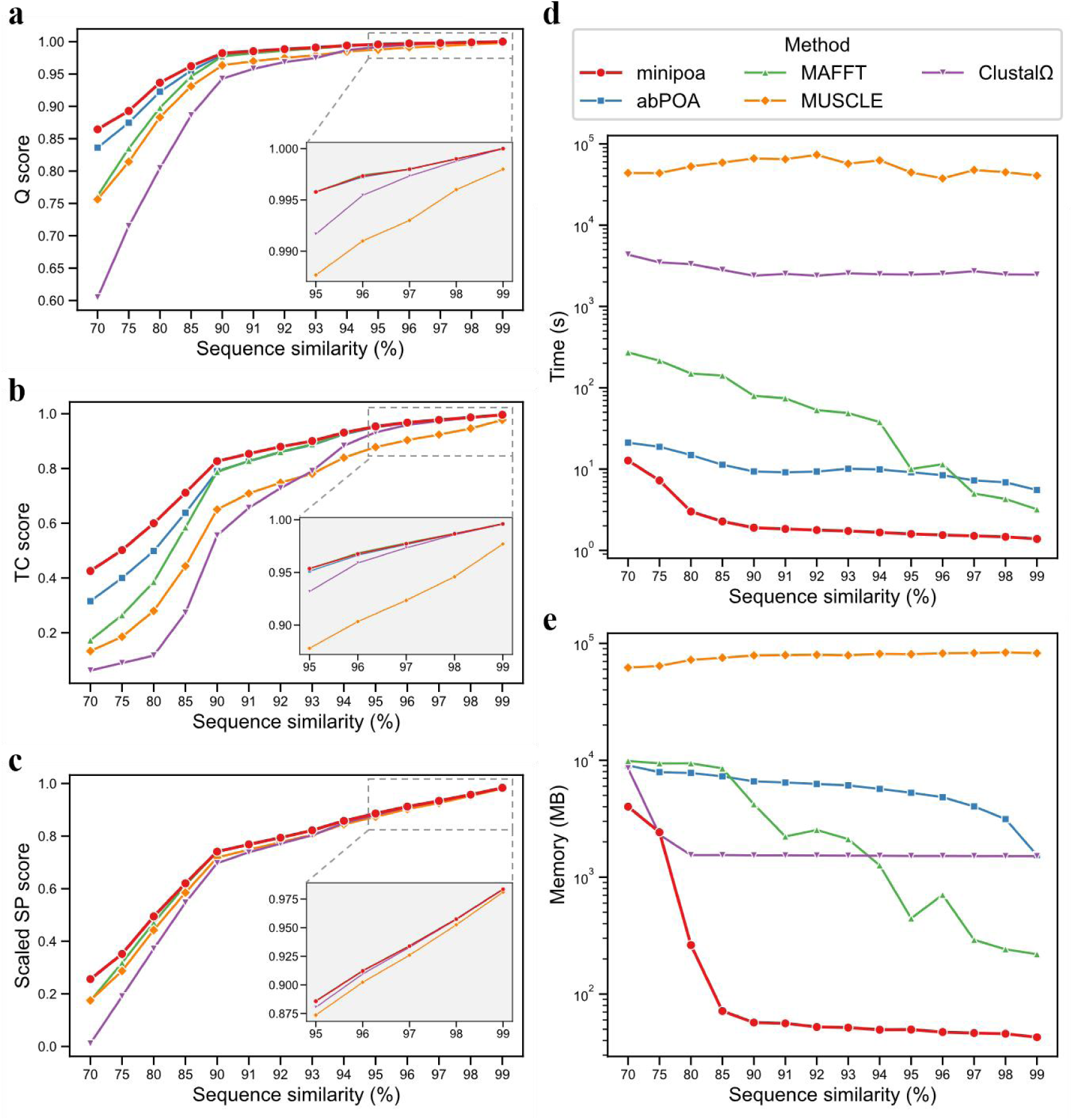
Comparison of MSA methods on simulated datasets across sequence similarity levels. (a) Q score. (b) TC score. (c) Scaled SP score. (d) Runtime. (e) Memory usage.

As shown in Figure 4, minipoa consistently achieved the highest alignment quality across all sequence similarity levels, substantially outperforming the classical tools. Among the other methods, abPOA and MAFFT generally delivered the next best results, while MUSCLE and ClustalΩ showed the lowest accuracy. Notably, under low-similarity conditions where most existing methods struggle, minipoa maintained remarkable performance. At the lowest similarity level of 70%, for example, it achieved a Q score of 0.865 and a TC score of 0.425. In contrast, all other tools scored below 0.84 in Q and below 0.32 in TC. This demonstrates that minipoa achieves superior accuracy and stability under low-similarity conditions, outperforming existing methods through the use of POA and optimized backtracking.

Minipoa consistently exhibits the best performance among the evaluated methods in terms of both computational efficiency and memory usage. Indeed, it achieves improvements in runtime and memory efficiency that are close to or even reach an order of magnitude on datasets with sequence similarity ≥ 80%, consistently outperforming all other methods. In contrast, MAFFT shows a clear increasing trend in both runtime and memory usage as sequence similarity increases. Although abPOA demonstrates relatively good alignment speed compared with other methods, its memory footprint is substantially larger. In summary, minipoa not only maintains excellent alignment accuracy but also delivers outstanding efficiency, with minimal time and memory overhead even on challenging datasets with low sequence similarity.

#### 2.3.2 Multiple Sequence Alignment on Real Data

We evaluated the MSA mode of minipoa on five genomic datasets spanning a range of lengths and similarity levels, and compared its performance with four widely used MSA tools. The datasets included mitochondrial genomes, SARS-CoV-2, HIV sequences, 16S rRNA, and MPox (see Supplemental Method).

As summarized in Table 3, only minipoa and MAFFT successfully completed alignments on all datasets. Other tools either failed on certain datasets or were terminated after three days without completion, which was considered a failure. For instance, abPOA did not finish the MPox alignment, ClustalΩ failed on MPox and required extremely long runtimes on most datasets, and MUSCLE did not complete any dataset. In terms of alignment accuracy, minipoa and abPOA both achieved excellent alignment accuracy on highly similar datasets such as mitochondrial and SARS-CoV-2 sequences. Notably, on lower-similarity datasets, minipoa substantially outperformed other aligners. For instance, on the HIV dataset, minipoa achieved an SP score of 0.20, whereas MAFFT and abPOA only reached 0.11. Similarly, on the moderately similar MPox dataset, minipoa delivered the highest alignment quality among all tested tools. Regarding computational efficiency, minipoa was the fastest across all datasets, often completing tasks nearly an order of magnitude faster than competing tools.

**Table 3.** Performance of MSA tools across five datasets.

| Method | *Mitochondrial genomes* |  | *SARS-CoV-2* |  | *HIV* |  | *16S rRNA* |  | *MPox* |  |
| --- | --- | --- | --- | --- | --- | --- | --- | --- | --- | --- |
|  | Time | Scaled SP | Time | Scaled SP | Time | Scaled SP | Time | Scaled SP | Time | Scaled SP |
| minipoa | 10s | 0.9914 | 11s | 0.9286 | 6min55s | 0.2023 | 13min56s | 0.0648 | 19min58s | 0.7346 |
| abPOA | 44s | 0.9913 | 2min20s | 0.9274 | 39min39s | 0.1146 | 38min17s | -0.0471 | — | — |
| MAFFT | 2min19s | 0.9910 | 25min54s | 0.8689 | 3h8min | 0.1078 | 3h49min | 0.0059 | 21h43min | 0.7025 |
| MUSCLE | >>72h | — | — | — | >>72h | — | — | — | — | — |
| ClustalΩ | 6h44min | 0.9913 | 11h43min | 0.4764 | 21h28min | 0.1993 | 15h2min | -0.4376 | >>72h | — |

Note: “—” indicates that the tool failed to complete the alignment. >>72h indicates that the tool was terminated after running for more than three days (72 hours).

### 2.4 Experimental Results on Large-scale Datasets

#### 2.4.1 Alignment of Million-base M. tuberculosis Genomes

We evaluated minipoa on large M. tuberculosis genomes of varying lengths from the POASTA benchmark[17]. Table 4 summarizes the runtime, memory usage, and alignment quality (Scaled SP) of MAFFT, POASTA, and minipoa on datasets of 250 kbp, 500 kbp, and 1 Mbp.

**Table 4.**
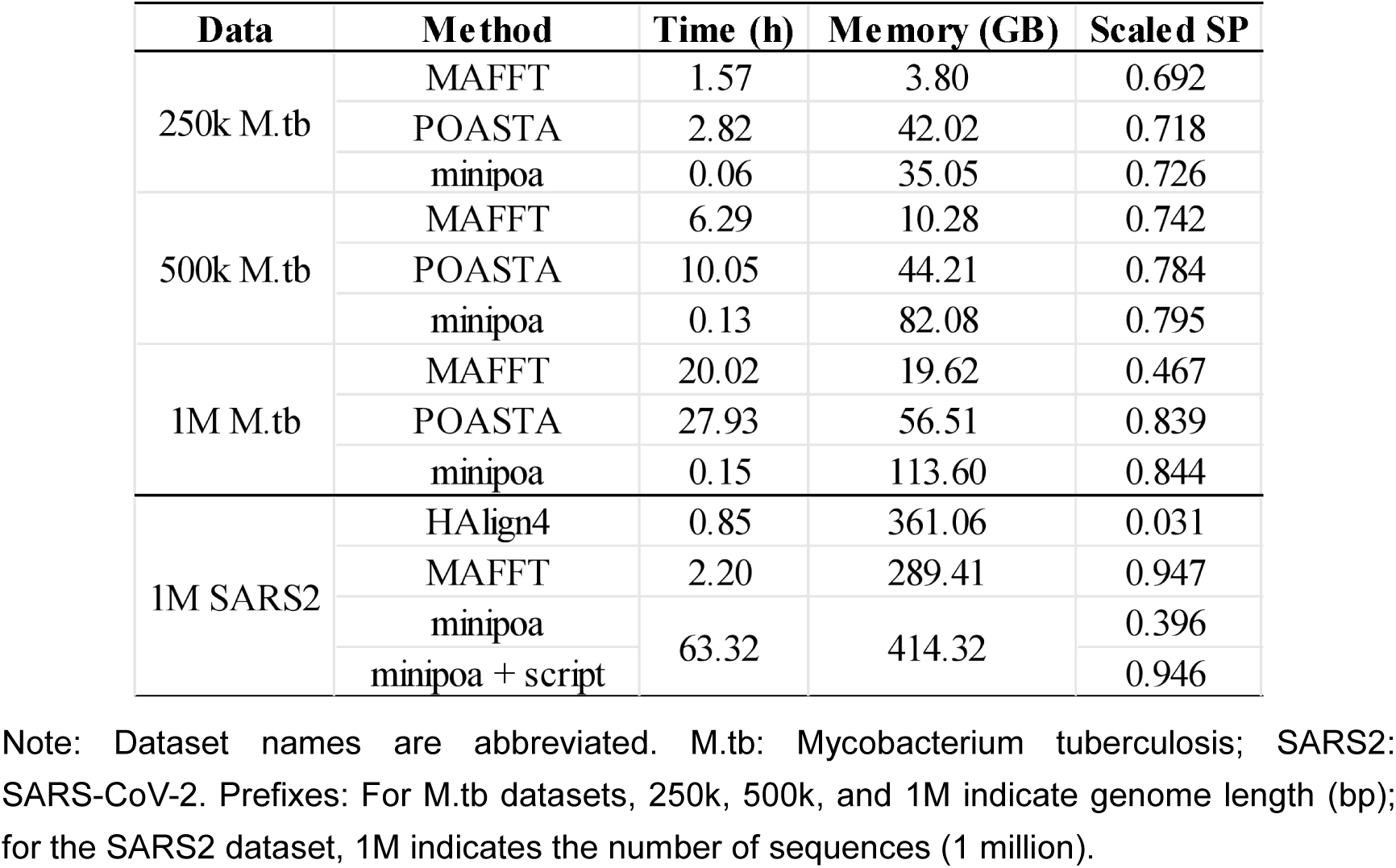
Performance of MSA tools on Large-scale Datasets.

In terms of alignment accuracy, minipoa consistently achieved the highest Scaled SP scores across all genome sizes. For the 250 kbp dataset, minipoa obtained a Scaled SP of 0.726, outperforming POASTA (0.718) and MAFFT (0.692). This trend was maintained for the 500 kbp dataset, where minipoa achieved 0.795, compared to 0.784 for POASTA and 0.742 for MAFFT. For the 1 Mbp dataset, minipoa reached 0.844, slightly higher than POASTA (0.839) and substantially higher than MAFFT (0.467), indicating that minipoa preserves high-quality alignments even as genome length increases. Notably, MAFFT’s alignment quality decreased substantially on the 1 Mbp dataset, highlighting the limitations of traditional MSA methods for very large genomes.

Regarding runtime efficiency, minipoa demonstrated a dramatic speed advantage. While MAFFT required over 20 hours to align the 1 Mbp genomes, and POASTA took nearly 28 hours, minipoa completed the same alignment in only 9 minutes. For memory usage, MAFFT used the least memory, while minipoa required the most. Both tools showed a roughly linear increase in memory consumption with genome length. POASTA required moderately high memory, which remained relatively stable regardless of dataset size. Overall, on high-similarity genomes, minipoa achieves a favorable balance between accuracy, efficiency, and scalability, making it highly practical for million-base genome alignments.

#### 2.4.2 Alignment of Million SARS-CoV-2 Sequences

The variants of concern of SARS-CoV-2 have highlighted the need for a global molecular surveillance of pathogens via whole genome sequencing. Such sequencing, for SARS-CoV-2 and other pathogens, is performed by an ever increasing number of labs across the globe, resulting in an increased need for an easy, fast, and decentralized analysis of initial data[28]. In the workflow of genomic epidemiology, MSA serves as the foundational prerequisite for downstream analyses, including variant calling, lineage assignment, and phylogenetic reconstruction.

At present, MAFFT and HAlign series[29–31] remain among the very few tools capable of performing multiple sequence alignment on SARS-CoV-2 datasets at the scale of millions of genomes. MAFFT, in its keeplength mode, is a progressive alignment method based on fast Fourier transform. In large-scale homologous sequence alignment, the keeplength mode ensures that the output alignment length is identical to the reference genome, facilitating downstream analyses based on a standard coordinate system. Its main advantage is speed and coordinate consistency, but a notable drawback is that insertions relative to the reference are forcibly removed, potentially losing biologically meaningful information. As a representative tool of the HAlign series, HAlign4[30] employs a center-star strategy and typically leverages suffix trees to rapidly identify shared substrings. It aligns all sequences to a reference sequence and merges the pairwise alignments into a global MSA. HAlign4 can handle extremely large datasets with relatively low memory usage while preserving all sequence information, including insertions, but its alignment quality may decrease when sequences exhibit high divergence.

For this study, we retrieved 17,231,704 publicly available SARS-CoV-2 genome sequences from the GISAID[32] database on June 15, 2025, with an average sequence length of 29,313.29 bp. From this collection, one million sequences were randomly sampled to construct a large-scale test dataset for performance evaluation. The standard reference genome, Wuhan-Hu-1 (Accession ID: NC_045512.2), was obtained from the NCBI GenBank[33] database.

We first evaluated the original minipoa alignment, in which insertions relative to the reference were retained, and compared its SP score with that produced by HAlign4. As shown in Table 4, minipoa achieved an SP score of 0.396, which is substantially higher than that of HAlign4 (0.031), indicating superior overall alignment quality when preserving full sequence information. Next, we evaluated the post-processed minipoa alignment after removing gap columns from the reference sequence and compared it with the MAFFT keeplength result. After reference-gap removal, minipoa achieved an SP score of 0.946, which is comparable to that of MAFFT (0.947). This close performance is noteworthy given the fundamental methodological differences between the two approaches. MAFFT in keeplength mode effectively aligns each query sequence to the reference through pairwise alignment, whereas minipoa performs MSA. As a result, a very small fraction of sequences in minipoa may not be optimally aligned to the reference sequence if a better match exists with other sequences in the dataset, leading to a marginal difference in SP score.

In terms of computational performance (Table 4), HAlign4 completed the alignment the fastest (0.85 hours) but used more memory (361.06 GB) than MAFFT. MAFFT was slightly slower (2.20 hours) but the most memory-efficient (289.41 GB). Minipoa required the most time and memory in its original form (63.32 hours, 414.32 GB) but produced much higher alignment quality. After reference-gap removal, minipoa achieved near-parity with MAFFT in SP score.

In summary, minipoa represents the only currently available tool for true multi-sequence alignment of SARS-CoV-2 genomes at the million-sequence scale, providing high-quality alignments that preserve biologically meaningful information. It can be applied directly to downstream analyses such as variant calling and phylogenetic reconstruction, offering a practical solution for large-scale genomic epidemiology.

## 3 Discussion

In this study, we present minipoa, a fast and memory-efficient partial order alignment software designed to address the growing demands of large-scale sequencing and multiple sequence alignment tasks. By combining a seed–chain–align paradigm, adaptive or static banded dynamic programming, and SIMD optimization, minipoa achieves a favorable balance among alignment accuracy, computational efficiency, and scalability across a wide range of sequence lengths and similarity levels.

A key strength of minipoa lies in its unified yet task-specific design. Sequencing and MSA tasks impose fundamentally different requirements: error correction typically operates on highly similar reads and benefits from aggressive pruning of the DP search space, whereas MSA must remain sensitive to substantial divergence among sequences. By explicitly separating sequencing and MSA into static-band and adaptive-band modes, minipoa avoids the inherent trade-offs between efficiency and sensitivity imposed by a single banding strategy. This design choice enables minipoa to retain high accuracy while significantly reducing memory usage, particularly in challenging cases involving long reads or low-similarity sequences.

In sequencing mode, the static-band design is particularly effective because the high similarity among reads strongly constrains the location of the optimal alignment path. By fixing a narrow DP band, minipoa limits unnecessary exploration of the alignment space and prevents excessive growth of DP states as the POA graph becomes larger. As a result, minipoa achieves consistently lower memory usage and faster runtimes than abPOA and TSTA in error-correction pipelines, while producing consensus sequences of comparable accuracy.

In MSA mode, the seed-chain-align strategy plays a central role in improving both speed and robustness. By performing seed collection and chaining on a linear consensus sequence rather than directly on the graph, minipoa transforms a complex graph-sequence matching problem into a more tractable sequence-sequence problem. The resulting anchor chains provide reliable global guidance for subsequent sequence-to-graph alignment, allowing the DP computation to focus on a narrow and informative region of the alignment space. This strategy is particularly effective for high-similarity datasets, where a well-distributed chain of anchors can often be identified. As a result, in high-similarity scenarios, minipoa is able to complete alignments that other tools fail to handle, such as MSAs with millions of sequences.

The improved backtracking strategy further enhances alignment accuracy in low-similarity regions. By incorporating graph edge weights into backtracking decisions, minipoa avoids spurious match paths introduced by weakly supported substitution nodes, a failure mode that is difficult to correct using standard DP backtracking rules. This graph-aware refinement can improve alignment consistency in simulated MSA benchmarks, leading to higher Q and TC scores under low-similarity conditions.

The error-correction experiments demonstrate that minipoa can be seamlessly integrated into established long-read correction pipelines as an efficient POA backend. When replacing the original SPOA component in Racon, minipoa achieved correction accuracy comparable to existing approaches while consistently reducing runtime and memory consumption across diverse datasets. These results indicate that optimized POA implementations such as minipoa can directly benefit high-throughput sequencing workflows, where computational efficiency is increasingly a limiting factor, and suggest that minipoa represents a viable alternative for consensus generation in large-scale long-read error correction.

More broadly, the multiple sequence alignment results reveal that minipoa maintains robust alignment quality while scaling efficiently to datasets with increasing size and sequence divergence. This combination of accuracy and scalability is particularly relevant in the context of pangenome research, where graph-based representations are becoming central to capturing population-level genomic variation. By enabling efficient construction and refinement of partial order alignment graphs from large and heterogeneous sequence collections, minipoa provides a scalable foundation for graph-based alignment and comparative analysis, and may facilitate downstream applications such as pangenome graph construction, variant-aware alignment, and haplotype-resolved analyses.

In addition, the SARS-CoV-2 case study illustrates a concrete advantage of minipoa in large-scale viral genomics. Current reference-guided alignment strategies often rely on MAFFT with the keeplength option, which requires manual removal of gaps in the reference sequence to preserve coordinate consistency. In contrast, minipoa produces high-quality alignments without the need to delete reference gaps, while achieving alignment accuracy comparable to or exceeding MAFFT even after gap removal. This property simplifies preprocessing, preserves biologically meaningful reference coordinates, and is particularly advantageous for large-scale viral surveillance efforts where tens or hundreds of thousands of genomes must be aligned in a consistent coordinate system.

Despite its strengths, minipoa has several limitations. Its runtime still increases with the size and complexity of the alignment graph, which can become a bottleneck for highly heterogeneous datasets with substantial structural variation. Moreover, alignments are currently performed in the order of sequences in the input FASTA file, and no algorithmic strategy has been implemented to determine an optimal sequence order that could improve efficiency or alignment quality.

## 4 Conclusion

We present minipoa, a fast and memory-efficient partial order alignment tool. Through a combination of algorithmic innovations and a task-specific design, it is suitable for large-scale sequencing and multiple sequence alignment. In error correction, minipoa produces consensus sequences comparable to existing methods while reducing runtime and memory usage. For multiple sequence alignment, it scales efficiently to large and heterogeneous datasets, as demonstrated by million-sequence SARS-CoV-2 MSAs, while maintaining robust alignment quality. We anticipate that minipoa will provide a practical and scalable solution for pangenome research, high-throughput sequence analysis, and consensus sequence generation workflows.

## 5 Methods

The workflow of minipoa is illustrated in Figure 1. Minipoa takes a set of sequences in FASTA format as input. Optionally, users may provide an existing POA graph in GFA format to initialize the alignment; otherwise, minipoa automatically constructs an initial graph using the first sequence in the FASTA file. Minipoa employs an iterative alignment strategy (Figure 1A). Starting from the initial graph, it first derives a consensus sequence from the graph to serve as a reference. This consensus is then used to find a chain of high-confidence anchors against the query sequence. The resulting anchor chain guides the subsequent sequence-to-graph DP process, which is performed using an adaptive banding strategy. As query sequences are progressively aligned and incorporated into the graph, minipoa can produce two primary outputs:(i) a consensus sequence or a MSA, both exported in FASTA format; or (ii) the final graph structure itself, exported in GFA format.

Minipoa provides two operating modes. The MSA mode employs a seed–chain–align strategy combined with adaptive banding and supports output in both FASTA and GFA formats. The sequencing mode only adopts a static banding strategy and outputs a consensus sequence. Both modes effectively eliminate the *O*(*n*^2^) space requirement imposed by the adaptive banding strategy in abPOA, substantially improving the scalability of the software.

### 5.1 Chaining

#### 5.1.1 Collecting seed

To efficiently identify reliable matches between the query sequence and the graph, we first collect minimizers from both the consensus sequence derived from the graph and the query sequence. Matching minimizers between the two sequences are then used to generate seeds, each representing a short exact match with defined coordinates in both sequences. By operating on the consensus sequence rather than directly on the graph, the problem of seed detection between a graph and a query sequence is effectively transformed into a seed-finding problem between two linear sequences. This transformation substantially reduces computational overhead, enabling rapid seed collection. Moreover, the resulting seeds exhibit high positional consistency, which facilitates the construction of high-confidence anchor chains for subsequent alignment.

#### 5.1.2 Finding the optimal local chains

Our method uses the same seed chaining strategy as implemented in minimap2[18].. Formally, an anchor is a 3-tuple (x, y, z), representing interval [x − w + 1, x] on the reference matching interval [y − w + 1, y] on the query. Given a list of anchors sorted by, let *f*(*i*) be the maximal chaining score up to the -th anchor in the list. *f*(*i*) can be computed by:

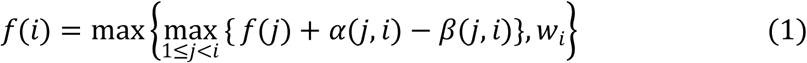

where α(*j,i*) = min{min {*y_i_* − *y_j_*, *x_i_*− *x_j_*}, *w_i_*} is the number of matching bases between anchor and . Let *d_ij_* = |(*y_i_* − *y_j_*,) –(*x_i_* − *x_j_*)| be the gap length. *β*(*j,i*) is the gap penalty:

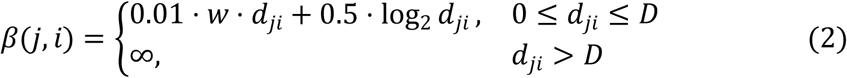

where *D* is 1000. Solving Eq. 1 leads to an *O*(n^2^) algorithm where is the number of anchors. This algorithm is extremely slow for large n. When evaluating Eq. 1, we start from anchor i − 1 and stop the process if we cannot find a better score after up to m iterations. This heuristic approach reduces the average time to *O*(*n* × *m*).

For the backtracking phase, we iterate through the list in reverse order. A backtracking procedure is initiated from each anchor that has not yet been marked as “visited” and all anchors belonging to the resulting path are subsequently marked as “visited”. This process yields several local chains, which may potentially intersect with one another.

#### 5.1.3 Finding the optimal chain

We define a local chain as a 5-tuple (t𝑠, t𝑒, q𝑠, q𝑒, A), where ts and te are the start and end positions in the reference sequence, qs and qe are the start and end positions in the query sequence, and A is the list of anchors within the chain. Given a list of local chains sorted by te, let *g*(*i*) be the maximal chaining score up to the i-th chain in the list. *g*(*i*) can be computed by:

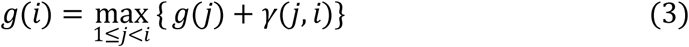

Here, γ(j, i) the incremental score for connecting the i-th chain to the j-th chain. Let n_i_ denote the number of anchors in the i-th chain, and f_i_(k) denote be the optimal score up to the k -th anchor in i-th chain’s own anchor list A_i_. The incremental score γ(j, i) is then given by:

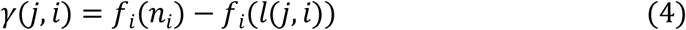

where *l*(*j*,*i*) is the smallest index in the i-th chain such that i-th chain is disjoint from the j-th chain after that anchor. Formally, l(j, i) satisfies:

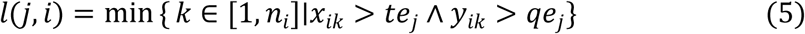

where *x_ik_* and *y_ik_* denote the x-value and y-value of the k-th anchor in the k-th local chain, respectively. Since the number of local chains is typically small (usually ≤ 25), we only need to accelerate the maximization step in Eq. 5 with a binary search. The chaining process is highly efficient and does not become a computational bottleneck in the overall workflow.

Following the backtracking step to identify the optimal chain, we perform a filtering procedure to refine the anchor set. Specifically, we iterate through the chain and selectively remove anchors to ensure that the distance between any two consecutive anchors is strictly greater than . In practice, we can almost always improve the result of alignment with 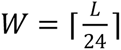, where *L* is the length of the reference and ⌈ ⋅ ⌉ denotes the ceiling function.

### 5.2 POA guided by anchor chain

The anchor chain is first projected onto the graph by mapping each consensus position to its corresponding graph nodes. Consecutive anchors partition the alignment into a series of local regions, within which the sequence-to-graph POA is performed. For each region, DP employs adaptive banding and SIMD optimizations, significantly speeding up the computation process while preserving alignment accuracy.

### 5.3 Band strategy in minipoa

Figure 5 illustrates the key differences between the k-band technique and the adaptive band strategy. In the k-band approach, the DP computation is restricted to a fixed band that is determined a priori, typically centered around the main diagonal inferred from a heuristic alignment. Once defined, this band remains unchanged throughout the DP process. In contrast, the adaptive band strategy dynamically adjusts the DP computation range based on information observed during the alignment process. Starting from an initial band, the algorithm decides whether to expand the band laterally or extend it downward according to intermediate DP states, allowing the computation region to better follow the true alignment path.

**Figure 5.**
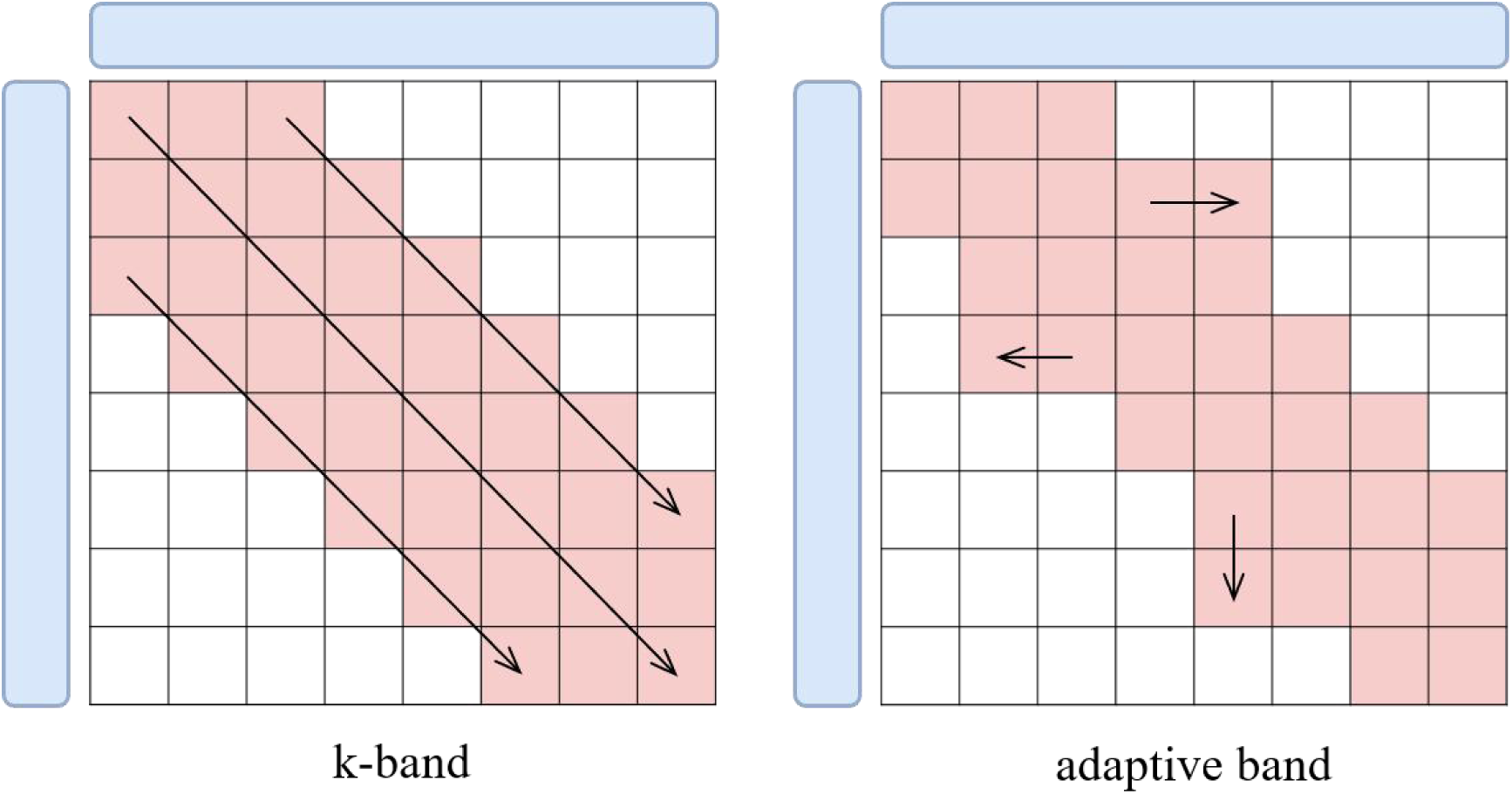
Comparison between the k-band and adaptive band strategies in DP alignment. In the k-band approach, the DP computation is restricted to a fixed band around the main diagonal, which is determined prior to the alignment and remains constant. In contrast, the adaptive band strategy dynamically adjusts the DP computation region based on intermediate DP states, expanding laterally or downward as needed to better follow the true alignment path.

#### 5.3.1 static band strategy

A static band strategy is employed for the sequencing mode, as this approach is optimal for datasets exhibiting high similarity. Conceptually, this approach can be viewed as analogous to the k-band technique in pairwise sequence alignment: a representative path in the graph is heuristically aligned to the query sequence, and dynamic programming is then restricted to a narrow region around the corresponding diagonal. The static banding process is only guided by a heuristic estimation of the best forward and backward path length for each node, calculated during a linear-time preprocessing stage (Algorithm 1).

**Algorithm 1:**
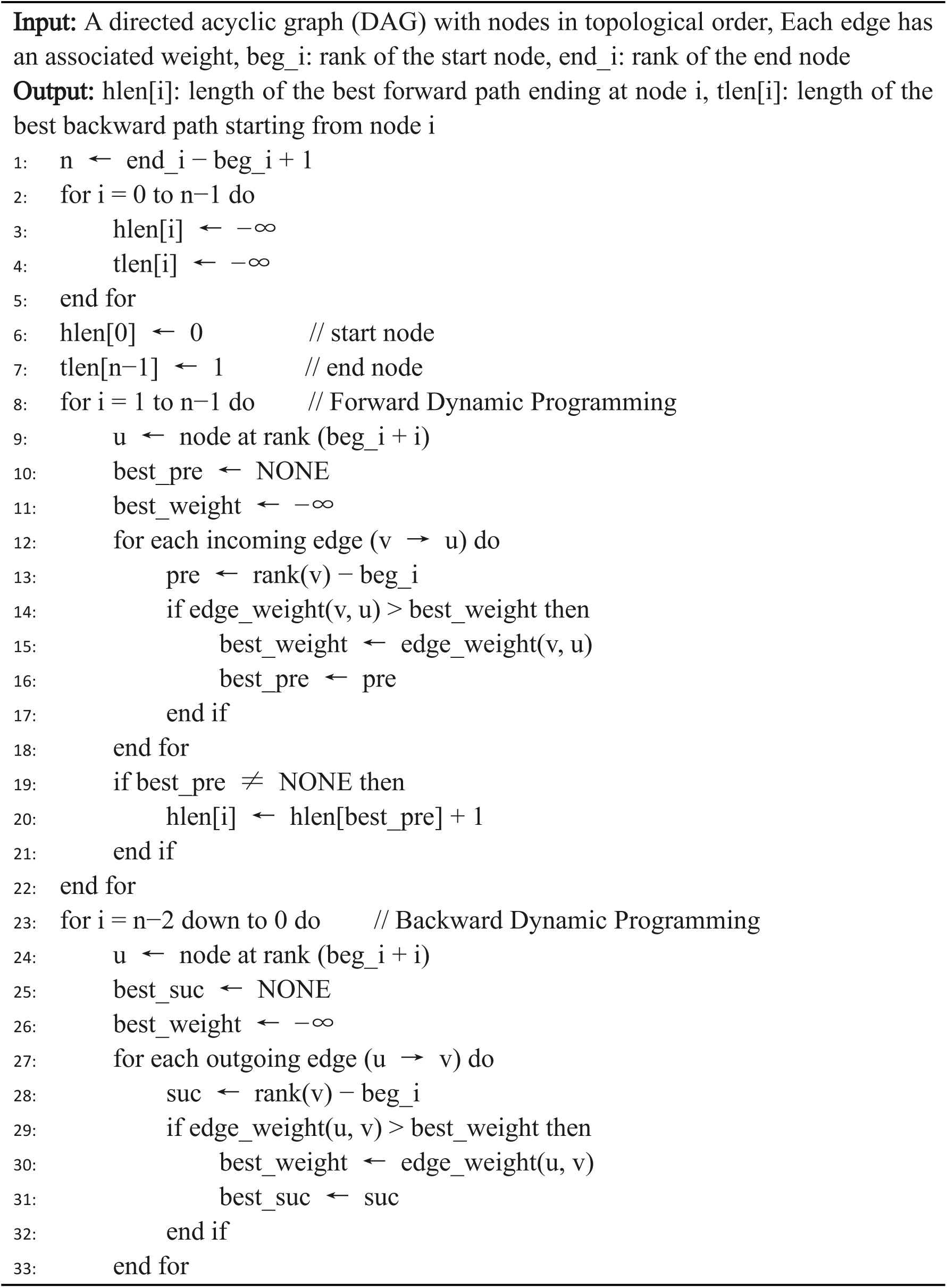
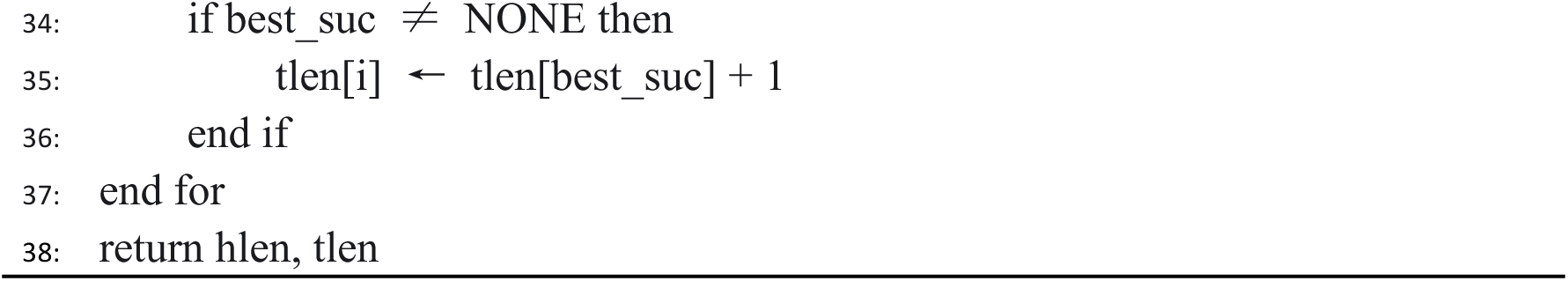
Computation of Forward Length and Backward Length

#### 5.3.2 adaptive band strategy

Our MSA mode enhances the adaptive banding technique from abPOA, enabling it to effectively handle datasets with a wide spectrum of sequence similarities. Our adaptive banding method is an extension of the static band strategy, the details of which are presented in Algorithm 2. For each node, the band is dynamically calculated based on several adaptive parameters. We track the length of the longest co-linear match path (Sl) ending at each node (lines 14-23). If no confident path is extended, the algorithm widens the band by increasing a penalty parameter Ow (lines 24-26), ensuring that the optimal path is not lost in regions of high divergence. If a high-confidence anchor path is identified (Sl[i] exceeds a threshold), we infer a reliable diagonal in the DP matrix. The algorithm then adjusts the band for subsequent nodes, centering it around this confident diagonal using the offset parameters Ol and Or (lines 27-31). These adaptive parameters (Pl, Pr, Ol, Or, Ow) are propagated forward through the graph to inform the calculations for all successor nodes (lines 32-36), allowing our method to be both fast in regions of high similarity and sensitive in regions of high divergence.

**Algorithm 2:**
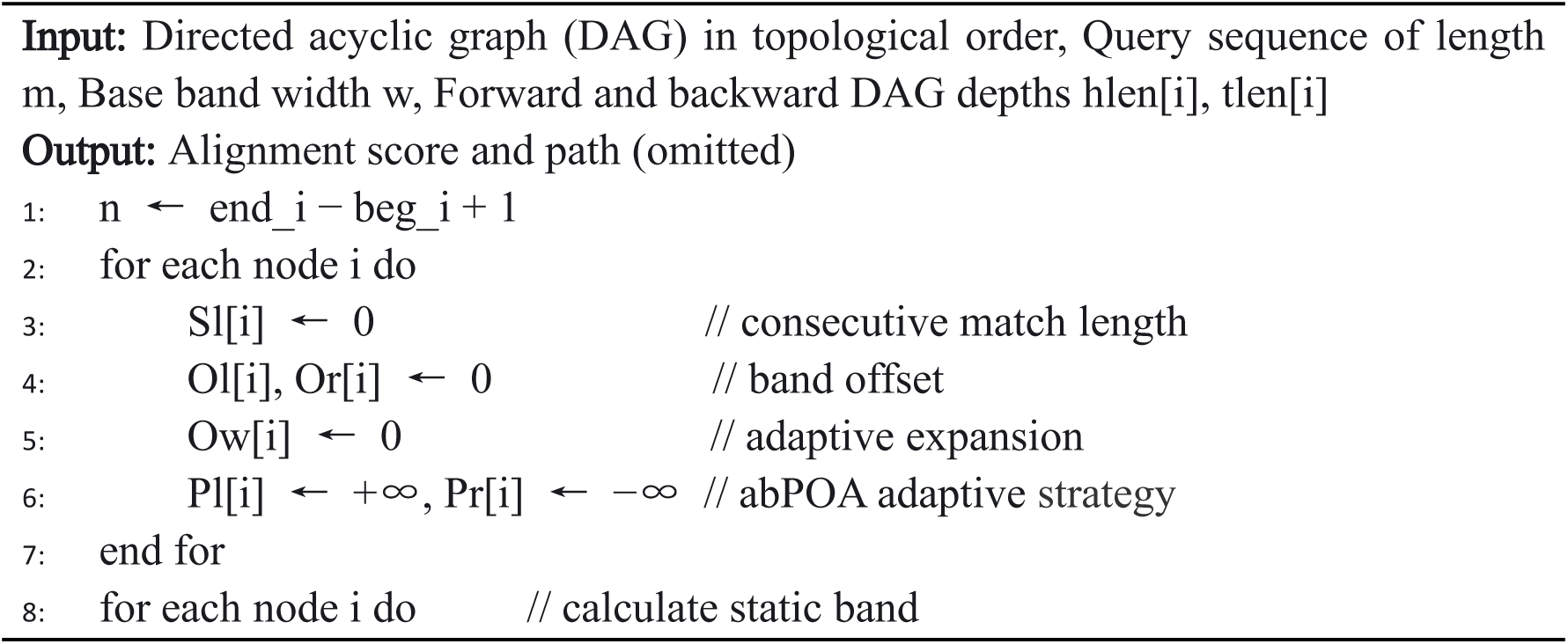
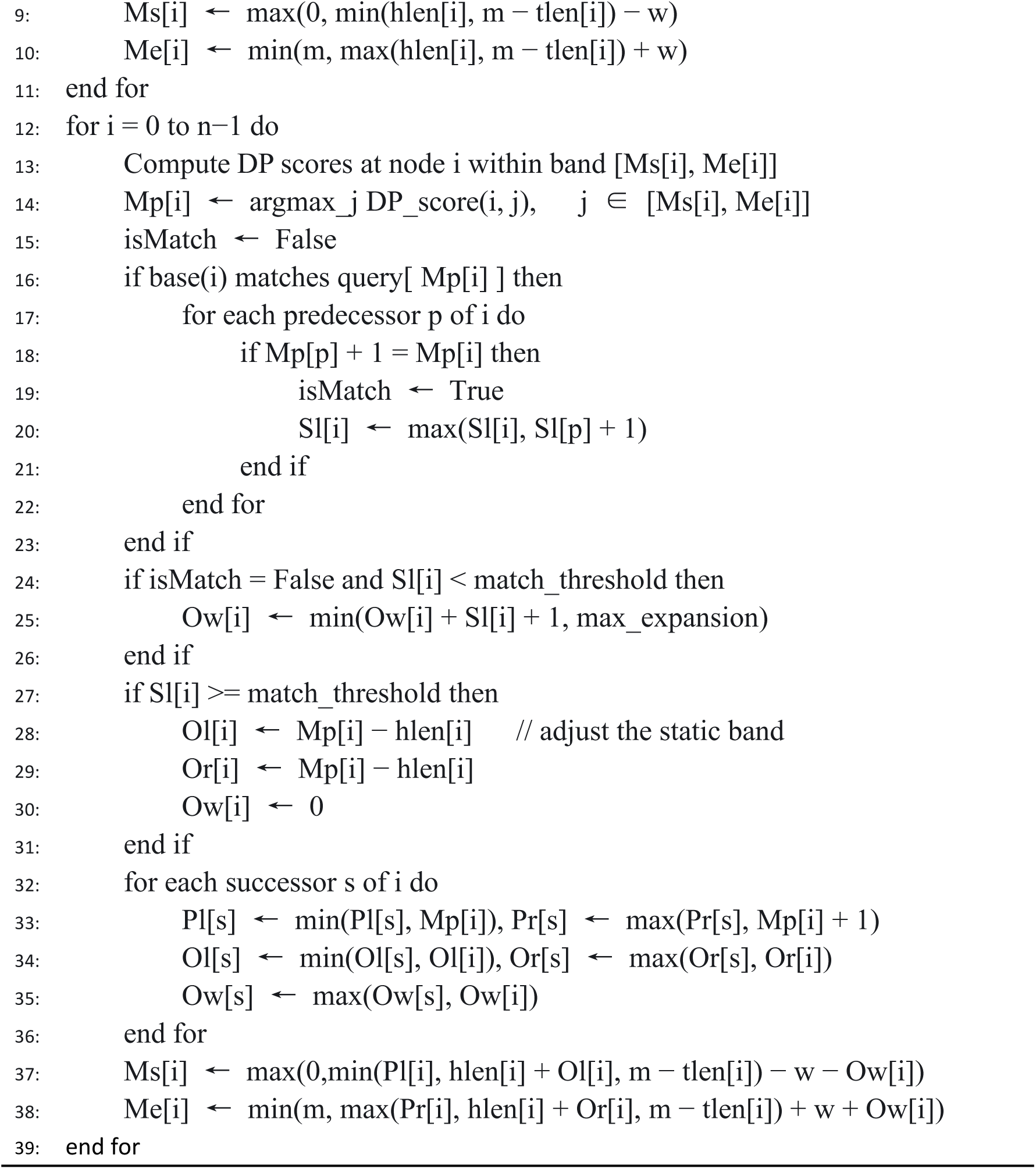
POA with Adaptive Banded Dynamic Programming

### 5.4 Optimization of Backtracking

Standard dynamic programming–based sequence alignment typically performs backtracking by following a fixed priority order, such as *match* → *deletion*→ *in𝑠𝑒rtion* → *mismatch*. While this strategy is effective for high-similarity alignments, it may lead to suboptimal or even incorrect alignments in low-similarity scenarios.

We propose a graph-aware backtracking strategy that leverages edge weights to prevent erroneous backtracking through low-confidence nodes. Specifically, during backtracking, we still follow the priority order (*m𝑎t𝑐ℎ_check_* → *del𝑒tion* → *in𝑠𝑒ction* → *match*→ *mismatch*). However, at the first occurrence of a match, we additionally examine whether the corresponding graph transition originates from a low-confidence node with insufficient edge support. If so, this match transition is rejected, and the backtracking process continues by considering deletion or insertion instead.

Figure 6 illustrates the difference between the conventional backtracking strategy and the proposed optimized approach. As shown in the bottom left, standard backtracking selects a match path that passes through a low-confidence node, leading to an incorrect alignment. In contrast, the optimized strategy (bottom right) identifies and avoids this transition, resulting in a more accurate alignment path.

**Figure 6.**
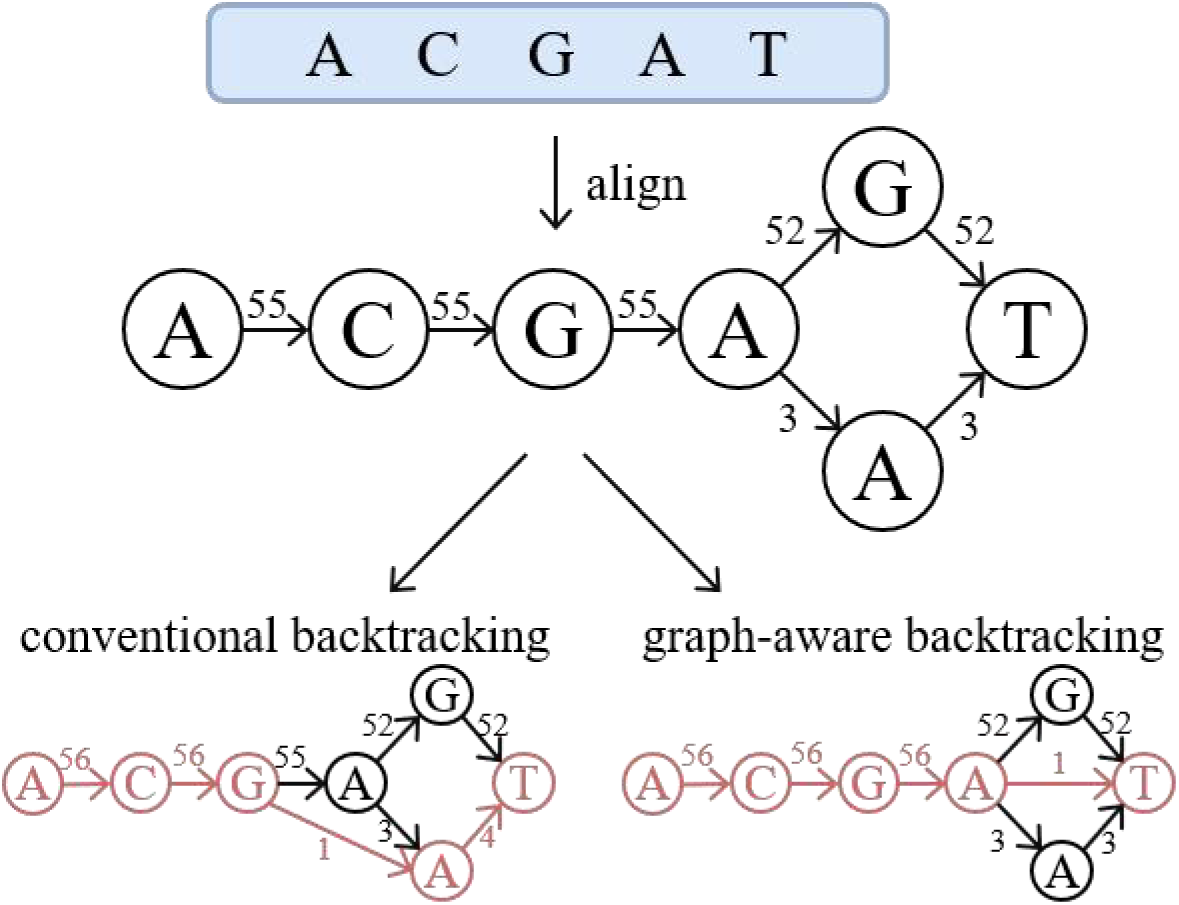
Comparison between conventional backtracking and the proposed graph-aware backtracking strategy. Conventional backtracking (bottom left) may select a path through a low-confidence node, resulting in an incorrect alignment. The graph-aware strategy (bottom right) avoids such transitions, yielding a more accurate alignment path.

## Acknowledgment

This work was supported by the National Natural Science Foundation of China (No. 62425107, No. 62473268) and Zhongguancun Academy, under the research projects Project No. 20240101 and Project No. 20240310. We sincerely appreciate their support in funding and resources.

## Software availability

The code of minipoa is located in https://github.com/NCl3-lhd/minipoa. The code of Racon-minipoa is located in https://github.com/NCl3-lhd/racon_minipoa. The version 1.0 of minipoa and Racon-minipoa can be found at Zenodo (https://zenodo.org/records/18676437).

## Data availability

The public datasets used in this study can be found at Zenodo (https://zenodo.org/records/18676437). The standard SARS-CoV-2 reference genome is available in the NCBI GenBank database under Accession ID NC_045512.2. The complete collection of SARS-CoV-2 genome sequences is available in the GISAID database.

