## Supplement Material 1 for "Minipoa: A minimizer-based method for fast and memory-efficient partial order alignment"

**Supplementary Materials:**

### Supplemental Method

#### Command-Line Usage

Minipoa is a command-line multiple sequence alignment (MSA) tool. The general usage is:

minipoa [options] <input.fasta>

The available command-line options are listed below:

-i, --inc_fp	Incrementally align sequences to an existing graph.

-M, --match (default: 2)	Match score.

-X, --mismatch (default: -4)	Mismatch penalty.

-O, --gap_open (default: -4)	Gap opening penalty.

-E, --gap_ext (default: -2)	Gap extension penalty.

-t, --thread (default: 1)	Number of threads.

-b, --band_b (default: 100)	band parameter for static banded alignment.

-f, --band_f (default: 40)	band parameter for static banded alignment.

-B, --ab_band	Enable adaptive banded alignment.

-S, --seeding	Enable minimizer-based seeding and anchoring.

-W, --poa_w (default: 10000)	Minimum distance between adjacent anchors.

-k, --k_mer (default: 19)	k-mer length used for minimizer seeding.

-w, --mm_w (default: 10)	Minimizer window size.

-p, --progressive_poa	Enable progressive POA mode.

-r, --result (default: 0)	Output format:

0 – consensus sequence

1 – row-column MSA format

2 – GFA graph format

-V, --verbose (default: 0)	Verbosity level:

0 – silent

1 – information

2 – debug

-h, --help	Print help message.

#### Running Modes of minipoa

Minipoa provides two specialized modes designed for different application scenarios.

1. Sequencing Mode

Sequencing mode is designed for long-read error correction and consensus generation. In this mode, minipoa outputs the consensus sequence by default.

Example: minipoa input.fasta > output.fasta

2. MSA Mode

MSA mode is designed for large-scale multiple sequence alignment tasks. In this mode, minipoa outputs the MSA format.

Example: minipoa input.fasta -S -r1 > output.fasta

#### *Simulation procedure and evaluation of read error correction on simulated data*

To simulate long-read datasets with varying read lengths and sequencing depths, we first randomly extracted a single 2mbp continuous reference sequence from the GRCh38 human reference genome.From this sequence, five subsequences with lengths of 500 bp, 5,000 bp, 20,000 bp, 50,000 bp, and 100,000 bp were further extracted and used as reference sequences for simulation. For each reference length, three sequencing depths (10×, 30×, and 50×) were considered, resulting in a total of 15 (5×3) simulation settings. For each setting, 100 independent clusters of sequences were generated from the same reference using PBSIM2[1], each representing an independent random simulation, with error profiles corresponding to Oxford Nanopore Technologies (ONT) or Pacific Biosciences (PacBio) sequencing.

We also modified the source code of PBSIM2 to only generate sequences coming from the forward strand. The modified source code of PBSIM2 is available at <https://github.com/malabz/pbsim2.>

To simulate PacBio data, PBSIM2 was run with the following arguments:

pbsim input.fasta --hmm_model data/P6C4.model --seed i --length-max length --length-min length --length-mean length --length-sd 0 --depth depth --accuracy-min 0.80 --accuracy-mean 0.85 --accuracy-max 0.90 --prefix pbsim_i

To simulate ONT data, PBSIM2 was run with the following arguments:

pbsim input.fasta --hmm_model data/R103.model --seed i --length-max length --length-min length --length-mean length --length-sd 0 --depth depth --accuracy-min 0.80 --accuracy-mean 0.85 --accuracy-max 0.90 --prefix ontsim_i

Three software packages were employed to generate consensus sequences: minipoa, abPOA[2], and TSTA[3]. The source code for these tools is publicly available at:

Minipoa: [https://github.com/NCl3-lhd/minipoa](https://github.com/NCl3-lhd/minipoa" \t "_new),

abPOA: [https://github.com/yangao07/abPOA](https://github.com/yangao07/abPOA" \t "_new),

TSTA: [https://github.com/bxskdh/TSTA](https://github.com/bxskdh/TSTA" \t "_new).

The run settings for each tool were as follows.

Minipoa:

minipoa input.fasta > output.fasta

abPOA:

abPOA input.fasta -M 2 -X 4 -O 4,24 -E 2,1 -r0 > output.fasta

TSTA:

TSTA-i input.fasta -m 2 -X -4 -T 30 -W 30 -o output.fasta

Alignment quality was evaluated by calculating the error rate of the generated consensus sequences. Specifically, each consensus sequence was aligned to its corresponding original reference sequence using Minimap2[4] (available at https://github.com/lh3/minimap2). The error rate was defined following the formulation in [2], as the total number of mismatches, insertions, and deletions in the alignment divided by the length of the consensus sequence.

Minimap2 was executed with the following settings:

minimap2 -a -x map-pb/map-ont --MD input.ref input.fasta > output.sam

#### Detail and evaluation of read error correction on real data

To evaluate minipoa under real-world usage scenarios, we modified the source code of the widely used long-read error correction pipeline, Racon (v1.4.13), replacing SPOA with minipoa in the multiple sequence alignment and consensus calling steps. The modified implementation, referred to as Racon-minipoa, is available at <https://github.com/NCl3-lhd/racon_minipoa.> We ran this modified Racon (Racon-minipoa), Racon-abPOA (available at https://github.com/yangao07/abPOA/releases/tag/v1.0.5) and the original Racon (available at https://github.com/lbcb-sci/racon/releases/download/1.4.13/racon-v1.4.13.tar.gz) in the error correction mode on five real datasets from different genomes (Table S1) and then calculated the error rates of the corrected reads. These datasets were originally used in the Racon paper to benchmark its performance[5].

The self-overlapping result of each input dataset was generated using minimap2 with the following settings:

minimap2 -t16 -x ava-ont/ava-pb input_seq.fa input_seq.fa > input_seq_ava.paf

The run settings of Racon-minipoa, Racon-abPOA and Racon-SPOA were identical:

racon input_seq.fa input_seq_ava.paf input_seq.fa -f -w 500 -t 16 > correct_seq.fa

The quality of corrected reads was assessed by mapping them to the corresponding reference genome using Minimap2. The error rate was calculated as the total number of mismatches, insertions, and deletions in the alignment divided by the length of the corrected read.

#### Detail and evaluation of MSA on real data

We evaluated the MSA mode of minipoa on five genomic datasets (Table S2) spanning a range of lengths and similarity levels, and compared its performance with four widely used MSA tools. The datasets included mitochondrial genomes, SARS-CoV-2, HIV sequences, 16S rRNA, and MPox. All datasets are available at [http://lab.malab.cn/%7Ecjt/MSA/datasets.html](http://lab.malab.cn/~cjt/MSA/datasets.html" \t "_new), except for the SARS-CoV-2 dataset. The SARS-CoV-2 dataset was constructed by randomly sampling 500 sequences from the Experiment of Million SARS-CoV-2 Sequences dataset. The sampling was performed without replacement, and no additional filtering or preprocessing was applied.

During multiple sequence alignment, all tools were executed using 8 threads, except abPOA, which only supports single-threaded execution. For all software, only the input and output files were specified, and default parameters were used, unless otherwise noted.

Minipoa was explicitly executed in MSA mode with the following settings:

minipoa input.fasta -S -W 500 -t 8 -r1 > output.fasta

Mafft was explicitly executed in AUTO mode with the following settings:

mafft --auto --thread 1 input.fasta > output.fasta

### Supplemental Tables

Table S1 Basic information of real datasets for read error correction

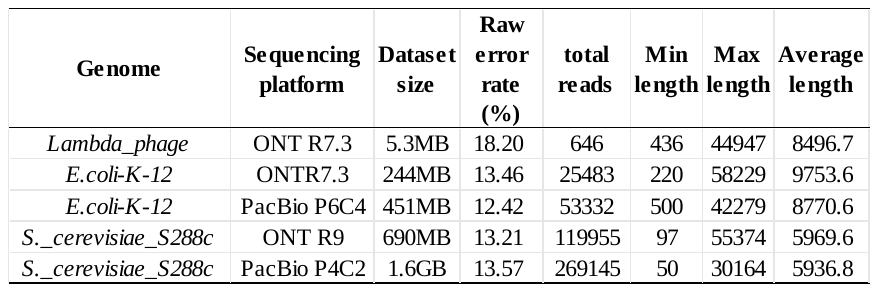

Table S2 Basic information of real datasets for MSA

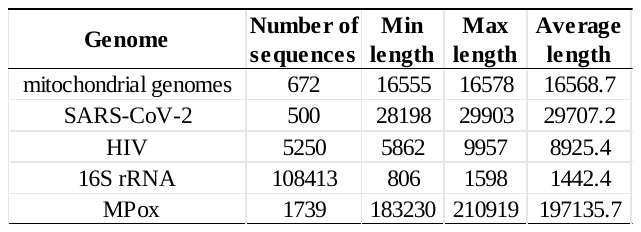
